# Elevated O-GlcNAcylation Enhances Pro-Inflammatory Th17 Function by Altering the Lipid Microenvironment

**DOI:** 10.1101/305722

**Authors:** Miranda Machacek, Zhen Zhang, Ee Phie Tan, Jibiao Li, Tiangang Li, Maria T. Villar, Antonio Artigues, Todd Lydic, Chad Slawson, Patrick Fields

## Abstract

Chronic, low-grade inflammation increases the risk of atherosclerosis, cancer, and autoimmunity in diseases like obesity and diabetes. Here, we show that increased levels of the nutrient-responsive, post-translational protein modification, O-GlcNAc (O-linked β-N-acetylglucosamine) are present in naïve CD4+ T cells from a diet-induced obesity murine model, and elevation in O-GlcNAc leads to increased pro-inflammatory IL-17A production. Importantly, CD4+ T helper 17 (Th17) cells, which secrete IL-17A, are increased in obesity and contribute to the inflammatory milieu. We found increased binding of the Th17 master transcription factor, RORγt, at the IL-17 locus and significant alterations in the lipid microenvironment, leading to increased ligands capable of increasing RORγt transcriptional activity. Importantly, the rate-limiting enzyme of fatty acid biosynthesis, acetyl CoA carboxylase 1 (ACC1), is necessary for production of these RORγt activating ligands and is O-GlcNAcylated. Thus, we have identified O-GlcNAc as a critical link between excess nutrients and pathological inflammation.

## INTRODUCTION

CD4+ T cells orchestrate the adaptive immune response. Activation of a naïve CD4+ T cell in response to a pathogen prompts differentiation of the naïve cell into various effectors (T helper 1 (Th1), Th2, Th17). Cytokines secreted by effectors then lead to elimination of the pathogen (Geginat et al., 2013). Importantly, major metabolic shifts occur during activation and are required for effector cell function. For example, activation induces a switch from oxidative phosphorylation to aerobic glycolysis (Chang et al., 2013; Frauwirth et al., 2002) and influx of glucose and glutamine is necessary to meet the energetic requirements for rapid clonal proliferation of the T cell (Macintyre and Divino Deoliveira, 2014; Swamy et al., 2016). Furthermore, different effectors require different metabolic pathways. For example, Th1, Th2, and Th17 cells utilize glycolytic pathways for energy, while regulatory T cells require oxidative phosphorylation (Michalek et al., 2011). Th17 cells have a requirement for endogenous fatty acid synthesis and pharmacological or genetic deletion of the rate-limiting enzyme in fatty acid biosysnthesis, acetyl CoA carboxylase 1 (ACC1), inhibits Th17 and favors regulatory T cell differentiation (Berod et al., 2014).

Significantly, metabolic abnormalities drive specific T cell effector pathology in several disease states. For example, the pro-inflammatory function of Th17 cells is enhanced in several autoimmune diseases, such as rheumatoid arthritis (RA) (Stadhouders et al., 2018). Recently, inflammatory Th17 cells infiltrating the synovium of joints in an RA model were found to have accumulation of lipid droplets due to increased fatty acid metabolism (Shen et al., 2017). Additionally, extrinsic metabolic factors alter T cell function. For example, in diseases of over-nutrition such as obesity and diabetes, Th1 and Th17 cells are increased in the peripheral blood and adipose tissue, contributing to atherosclerotic plaque formation and insulin resistance (Chehimi et al., 2017; Kanneganti and Dixit, 2012; Schindler et al., 2017; Winer et al., 2009). However, mechanisms that clearly link excess nutrients with aberrant T cell function are unclear.

The post-translational protein modification O-GlcNAc (O-linked β-N-acetylglucosamine) is an intriguing candidate for a how a T cell quickly interprets changes in nutrients. O-GlcNAc is added to nuclear, cytoplasmic, and mitochondrial proteins by a single enzyme, O-GlcNAc transferase (OGT) and removed by O-GlcNAcase (OGA) (Hart et al., 2011). O-GlcNAc levels are exquisitely sensitive to the nutritional and environmental status of the cell. The substrate for OGT, UDP-GlcNAc, is made through the hexosamine biosynthetic pathway and requires input from all four major biomolecules: carbohydrate, amino acid, fatty acid, and nucleic acid (Slawson et al., 2010). As nutrient flux changes through the cell, UDP-GlcNAc concentrations and O-GlcNAcylation by OGT change accordingly (Kreppel and Hart, 1999). Thus, in response to major nutrient shifts, rapid changes in O-GlcNAcylation of proteins and cellular function occur, suggesting that O-GlcNAcylation acts as a nutrient sensor to fine-tune cellular functions. O-GlcNAcylation is essential for murine and human T cell activation (Golks et al., 2007; Lund et al., 2016), a signaling event that triggers major metabolic changes. Intriguingly, abnormal levels of the O-GlcNAc processing enzymes contribute to T cell mediated diseases. For example, hypomethylation of the X-linked OGT gene in female lupus patients increases OGT expression and contributes to increased inflammatory T cells (Hewagama et al., 2013), while a miRNA targeting OGT is decreased in T cells of multiple sclerosis patients (Liu et al., 2017). Additionally, a single nucleotide polymorphism leading to truncated, nonfunctional OGA in a Mexican American population contributes to the increased incidence of diabetes in this population (Lehman et al., 2005). Thus, O-GlcNAc is essential for normal T cell function and aberrant O-GlcNAcylation is linked to inflammation.

In this study, we demonstrate that elevated O-GlcNAc levels increase pro-inflammatory IL-17A cytokine secretion from murine and human CD4+ T cells. We show a critical role of O-GlcNAc nutrient-sensing in regulating Th17 cell function, particularly in diet-induced obesity, and identify metabolic shifts in the lipid microenvironment that result in ligands capable of increasing IL-17A expression. Our results identify the O-GlcNAc modification as a critical mechanism for how Th17 cells translate nutrient excess into amplified pathological inflammation.

## RESULTS

### In murine CD4+ T cells, elevated O-GlcNAcylation increases pro-inflammatory IL-17A transcript and protein levels

To determine if O-GlcNAc plays a role in the function of T cell effectors, we treated murine splenic CD4+ T cells with Thiamet-G (TMG), a highly specific OGA inhibitor (Yuzwa et al., 2008), for 6 hours before activation under non-polarizing conditions (Th0). TMG treatment led to elevated O-GlcNAc levels over the four days of cell culture (Figure 1A). As has been previously observed (Hart et al., 2011; Zhang et al., 2014), levels of OGT decrease and levels of OGA increase in response to elevated O-GlcNAcylation in an attempt to restore steady state levels of O-GlcNAc. This is a well-described compensatory phenomenon, suggesting cells have a particular level of O-GlcNAcylation that is ideal for their function. Interestingly, re-stimulation of CD4+ T cells after 4 days of culture appears to “reset” or promote a restoration of the homeostatic level of O-GlcNAc, OGT, and OGA—a phenomenon which confirms previous reports (Golks, 2007; Hart, 1991). After activated CD4+ T cells differentiated over the course of four days, cells were re-stimulated and supernatants were harvested 24 hours later. The levels of key cytokines produced by four main effector types (Th1, Th2, Th17, and Treg) were analyzed by ELISA. Strikingly, IL-17A, the major cytokine secreted by Th17 cells, doubled (Figure 1B). Additionally, IFNγ, the major cytokine secreted by Th1 cells was also significantly increased. However, IL-4 and IL-10, cytokines predominantly produced by the Th2 and Treg lineages respectively, showed no significant changes. Since both IL-17A and IFNγ are pro-inflammatory cytokines and IL-4 and IL-10 have immunomodulatory functions, this suggests that elevated O-GlcNAcylation promotes increased inflammatory function of CD4+ T cells. Transcript levels of IL-17A, IFNγ, IL-4, and IL-10 followed similar trends to protein levels (Figure 1C). Additionally, another critical Th17 marker, IL-23 receptor (IL-23R), was also significantly increased. IL-23 binding and signaling through IL-23R stabilizes the Th17 phenotype (Aggarwal et al., 2003; Chen et al., 2007) and is also a marker of pathogenic Th17 function (Ghoreschi et al., 2010). Overall, markers of Th17 function are increased with elevated O-GlcNAcylation.

**Figure 1.**
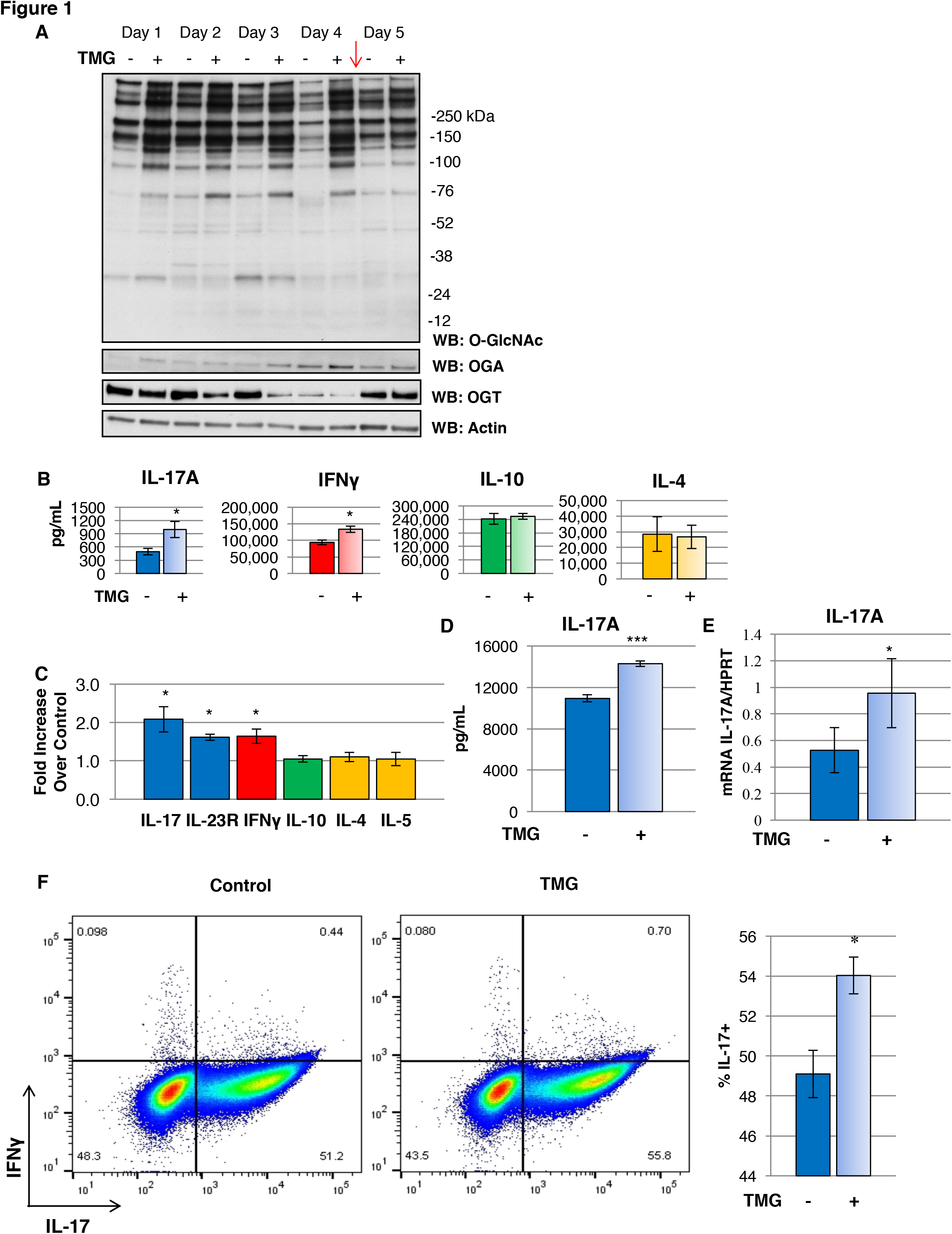
In murine CD4+ T cells, elevated O-GlcNAcylation increases pro-inflammatory IL-17A transcript and protein levels. A. TMG treatment increases O-GlcNAc levels over the course of CD4+ T cell differentiation and proliferation with corresponding decreases in O-GlcNAc transferase (OGT) and increases in O-GlcNAcase (OGA). The red arrow indicates the time of re-stimulation. B. Protein levels of cytokines secreted from CD4+ T cells with and without TMG treatment. C. Transcript levels of cytokines secreted from CD4+ T cells with and without TMG treatment. D. Protein level of IL-17A secreted by naïve cells polarized to Th17 lineage treated with and without TMG. E. Transcript level of IL-17A secreted by naïve cells polarized to Th17 lineage treated with and without TMG. F. Representative histogram showing CD4+ IL-17+ cells are significantly increased with TMG treatment. 25,000 cells plotted per condition. Dead cells were excluded. Bars represent mean +/− SEM of 5 biological replicates in B. and C. and 3 different biological replicates in D.-F.; * p < 0.05, *** p < 0.001

Th17 cells compose less than 1% of CD4+ T cells in the peripheral blood (Shen et al., 2009). To investigate the mechanism of O-GlcNAc regulation of Th17 function, we sought to obtain a more homogenous population of Th17 cells. Thus, we isolated murine splenic naïve CD4+ T cells (CD62L^hi^CD44^lo^) and treated with and without TMG before activation and polarization towards the Th17 lineage. Even with polarization towards a Th17 lineage, TMG treatment still significantly increased both protein and transcript IL-17A levels (Figure 1D and 1E). Intracellular cytokine staining showed a significant but modest increase in CD4+IL-17+ cells, suggesting there may be some effect on differentiation as well as functional cytokine output (Figure 1F).

### In human CD4+ T cells, elevated O-GlcNAcylation increases pro-inflammatory IL-17A transcript and cytokine levels

Th17 cells infiltrate adipose tissue to a greater extent in obese humans (McLaughlin et al., 2014) and secrete more IL-17A, contributing to adipose insulin resistance (Eljaafari et al., 2015). Thus, we wanted to determine if TMG treatment increased IL-17A from human CD4+ T cells. We obtained fresh whole blood from human volunteers through the Biospecimen Repository Core Facility (BRCF) at the University of Kansas Medical Center. Characteristics of volunteers, including gender, age, and ethnicity, are detailed in Figure 2A. CD4+ T cells were isolated from blood and treated with and without TMG before activation under Th0 and Th17 conditions. IL-17A significantly increased with elevated O-GlcNAc levels under both Th0 and Th17 polarizing conditions, recapitulating the results seen in mice (Figure 2B). Next we compared transcript levels of IL-17A and other pro-inflammatory cytokine markers, such as IL-23R and IFNγ, with and without TMG treatment in the CD4+ T cell population polarized to Th17 lineage. IL-17A and IL-23R, both transcriptional targets of RORγt, and IFNγ, the main pro-inflammatory cytokine from Th1 cells, were significantly increased with elevated O-GlcNAcylation (Figure 2C). Thus, we have demonstrated a clear effect of elevated O-GlcNAc levels on regulating pro-inflammatory cytokine signaling—particularly IL-17A—the major cytokine that mediates the function of Th17 cells.

**Figure 2.**
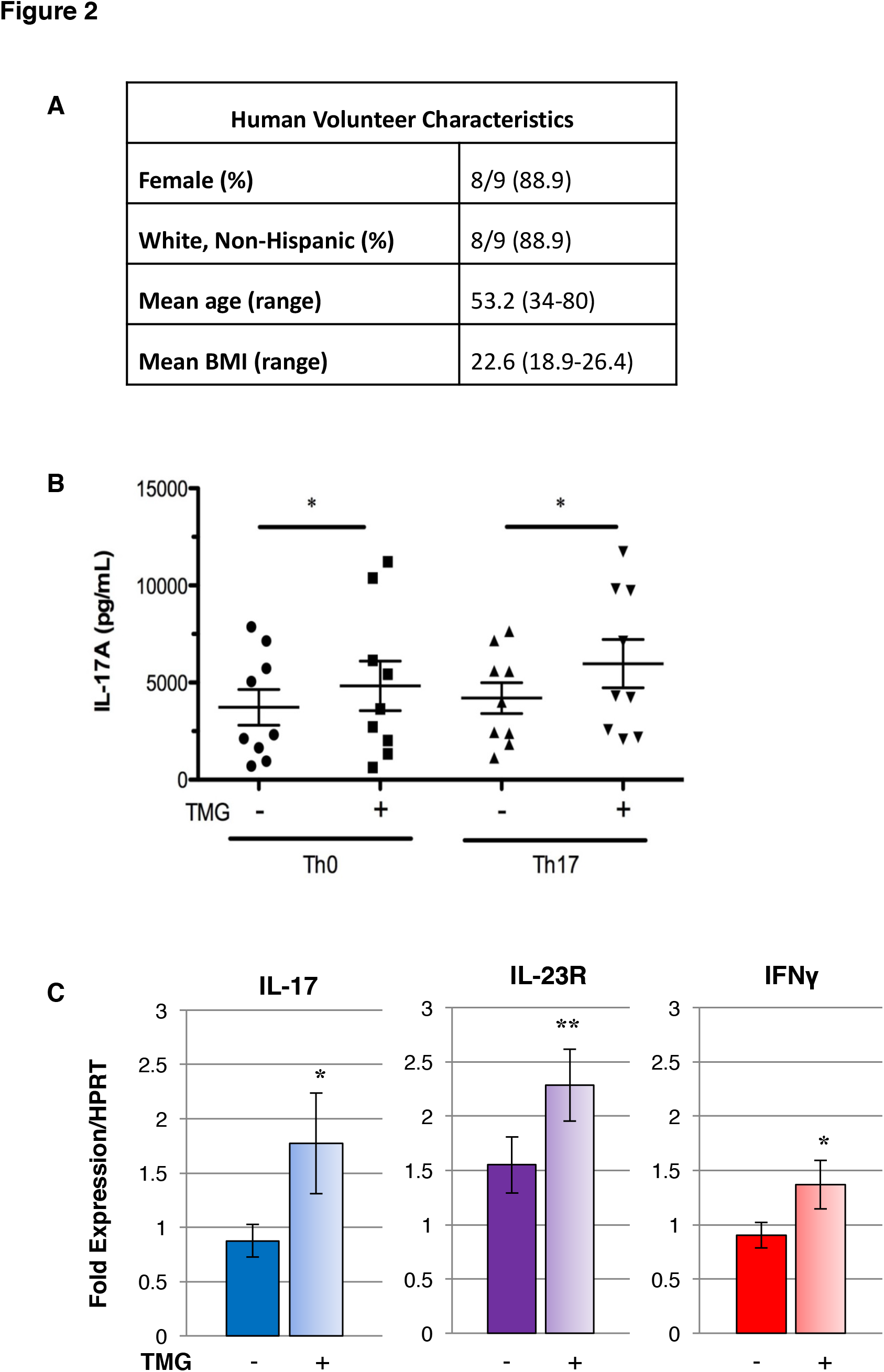
In human CD4+ T cells, elevated O-GlcNAcylation increases pro-inflammatory IL-17A transcript and cytokine levels. A. Clinical characteristics of human blood donors. B. TMG treatment increases IL-17A secretion from human CD4+ cells polarized under Th0 and Th17 conditions. C. IL-17, IL-23R, and IFNγ transcript levels all significantly increase with TMG treatment. Bars represent mean +/− SEM of 9 and 7 human donors in B. and C. respectively; * p < 0.05, **p < 0.01

### Naïve CD4+ T cells from Western diet fed mice have elevated O-GlcNAc levels and increased IL-17A production which is exacerbated by OGA inhibition

The previous data provide evidence that elevated O-GlcNAc increases pro-inflammatory cytokine production from CD4+ T cells. To investigate the physiological relevance of this effect in a disease model, we utilized a diet-induced obese mouse model (DIO). O-GlcNAc levels are sensitive to the nutritional state of the host and thus we would expect them to be elevated in a DIO model, phenocopying the effects of TMG in our *in vitro* studies. To test this hypothesis, we fed C57BL/6 male mice a high fat and cholesterol “Western” diet (WD) chow for 16 weeks. As expected, WD fed mice gained significantly more weight and their blood glucose was significantly elevated 15 weeks after initiation of the diet compared to mice fed standard chow (SC) (Figure 3A and 3B). Naïve CD4+ T cells (CD62L^hi^CD44^lQ^) isolated from WD fed mice had significantly elevated O-GlcNAc levels compared to SC fed mice (Figure 3C). Interestingly, OGT and OGA levels were not different between WD and SC fed mice, suggesting that elevated O-GlcNAc levels in the WD fed mice reach a new, albeit elevated and potentially pathogenic “set point.” When naïve CD4+ T cells from WD and SC fed mice were activated and polarized towards a Th17 lineage, cells from WD fed mice secreted more IL-17A as would be expected from previous studies (Chehimi et al., 2017; Winer et al., 2009). Strikingly, when the same naïve CD4+ T cells from WD and SC fed mice were treated with TMG before activation and polarization, cells from SC fed mice secreted levels of IL-17A comparable to cells from WD fed mice and IL-17A secretion from cells from WD fed mice was even further exacerbated (Figure 3D). The trends seen with IL-17A transcript levels were similar to the trends seen in protein levels (Figure 3E). Taken together, these results demonstrate that in a type 2 diabetes model, elevated O-GlcNAc levels in CD4+ T cells lead to a phenotype similar to pharmacological OGA inhibition by TMG. This suggests that O-GlcNAc is a critical regulator of Th17 functional activity and when aberrantly elevated—as in obesity—could contribute to the inflammatory milieu that drives the devastating pathology subsequent to type 2 diabetes.

**Figure 3.**
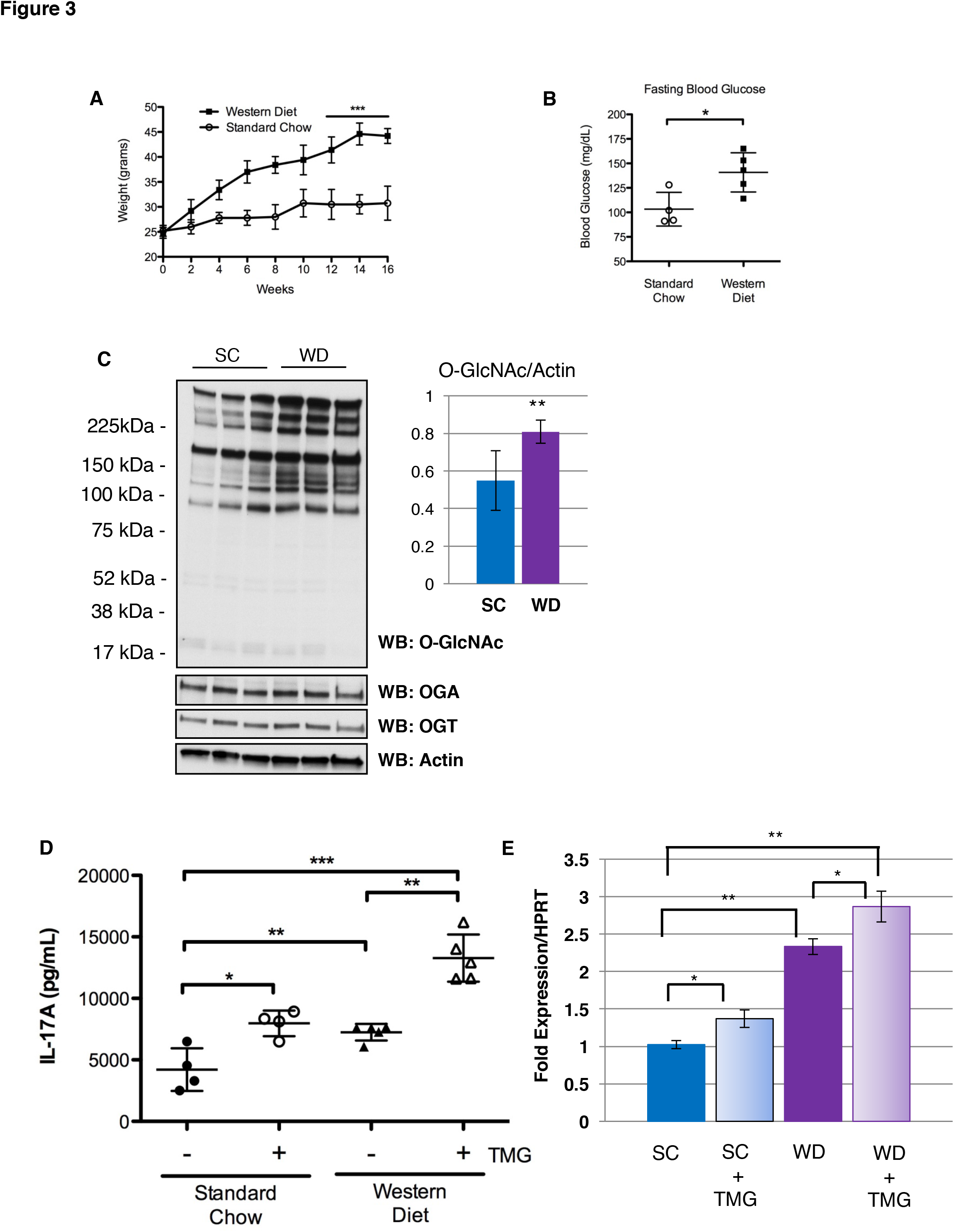
Western diet feeding in mice results in elevated O-GlcNAc levels in naïve CD4+ T cells and increases IL-17A secretion from naïve CD4+ T cells, which is exacerbated by TMG treatment. A. WD feeding results in significantly increased weight gain. B. WD feeding results in significantly elevated fasting blood glucose levels. C. WD fed mice have significantly elevated O-GlcNAc levels. D. Naïve CD4+ T cells polarized to the Th17 lineage from WD fed mice secrete significantly more IL-17A than cells from SC mice. Cells from SC fed mice treated with TMG secrete levels similar to cells from WD mice, and TMG treatment of cells from WD fed mice significantly exacerbates IL-17A secretion. E. IL-17A transcript levels mimic the trends seen with IL-17A protein secretion. In A. and B., points represent average −/+ SD and are calculated from 4 and 5 biological replicates (SC and WD respectively). In C., inset of densitometry is from 8 biological replicates and bars represent mean −/+ SD. Each lane in blot represents cell lysate from one mouse. In D. and E. bars represent mean −/+ SEM of 4 and 5 biological replicates (SC and WD respectively). HPRT is used an internal reference for gene expression. * p < 0.05, **p < 0.01, ***p < 0.001

### Elevated O-GlcNAcylation has no effect on RORγt protein or transcript levels but does promote retention of RORγt at the IL-17 locus

To investigate a mechanism for increased IL-17A secretion with OGA inhibition, we hypothesized that RORγt (retinoic acid related orphan receptor, t splice variant) would be increased with elevated O-GlcNAc levels. RORγt is the master transcription factor that directs the Th17 lineage and is essential for IL-17A gene transcription (Ivanov et al., 2006). Interestingly, we found no differences in RORγt by either protein or transcript level in the presence of TMG on the fourth day of cell culture (indicated as “0” time point) or over a 24 hour span after re-stimulation (Figure 4A and 4B). Furthermore, we saw no significant differences in RORγt protein or transcript levels at 24 hours after re-stimulation in Th17 differentiated cells from mice fed SC or WD (Figure S1A and S1B). These results suggest that O-GlcNAc neither regulates the stability of RORγt or its transcripts nor does O-GlcNAc affect transcription factors required for transcription of RORγt itself.

**Figure 4.**
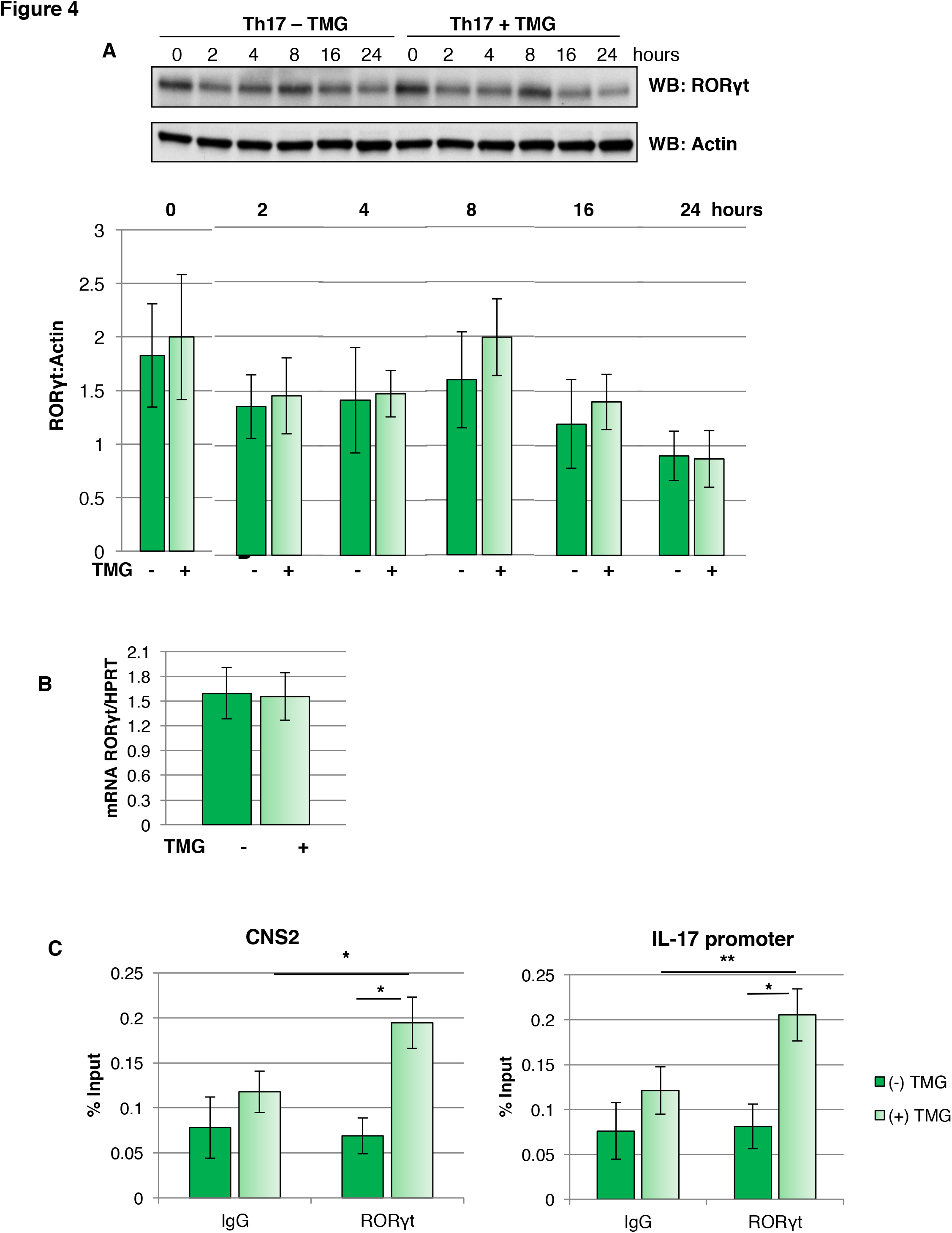
Elevated O-GlcNAcylation has no effect on RORγt protein or transcript levels but does promote retention of RORγt at the IL-17 locus. A. Prior to re-stimulation on the fourth day of culture (0) and over 24 hours after re-stimulation, RORγt protein levels are unchanged. Densitometry was used to compare RORγt to actin ratio at specified time points. Blot is representative of 3 biological replicates. Densitometry bars represent the mean +/− SEM of 3 biological replicates. B. Transcript levels of RORγt are unchanged 24 hours after re-stimulation. RORγt gene expression was normalized to HPRT expression. Bars represent mean +/− SEM of 5 biological replicates. C. TMG treatment significantly increases RORγt binding at the IL-17 promoter and CNS2 enhancer regions. Bars represent mean +/− SEM of 4 biological replicates; * p < 0.05, **p < 0.01

Since RORγt levels did not change with TMG treatment, we speculated that RORγt is being retained at the IL-17A locus. We performed chromatin immunoprecipitation (ChIP) of RORγt at the IL-17 promoter and an enhancer, conserved non-coding sequence 2 (CNS-2), which is required for IL-17A transcription (Wang et al., 2012). TMG treatment resulted in significantly increased RORγt binding at IL-17 promoter and the CNS-2 enhancer region in Th17 cells differentiated *ex vivo* and fixed on the fourth day of cell culture (Figure 4C). Taken together, TMG treatment promotes “locking in” of RORγt at the IL-17A promoter and enhancer regions.

### In murine differentiated Th17 cells, elevated O-GlcNAc increases lipid ligands capable of increasing RORγt transcriptional activity

Among the lineage defining transcription factors for CD4+ T cells, RORγt activity is uniquely regulated by the binding of lipid ligands. We thus investigated alterations in the lipidome that might increase RORγt transcriptional activity at the IL-17 gene. Cholesterol biosynthetic intermediates, such as 7β, 27-dihydroxycholesterol (7β, 27-OHC) and 7α, 27- dihydroxycholesterol (7α, 27-OHC), are known endogenous ligands capable of binding the ligand binding domain of RORγt, strongly activating its transcriptional activity (Santori et al., 2015; Soroosh et al., 2014). Additionally, fatty acids are critical for optimal RORγt activity as evidenced by a lack of Th17 differentiation under conditions of pharmacological inhibition or genetic deletion of the rate-limiting enzyme in fatty acid biosynthesis, acetyl CoA carboxylase 1 (ACC1) (Berod et al., 2014; Endo et al., 2015). The type of fatty acid ligand is also important for RORγt activity. An increased ratio of saturated fatty acids (SFA) to polyunsaturated fatty acids (PUFA) increases RORγt transcriptional activity at the pro-inflammatory IL-17A promoter while decreasing activity at the anti-inflammatory IL-10 promoter, thus promoting pathogenic Th17 differentiation (Wang et al., 2015). To investigate how O-GlcNAc alters the lipidome of T cells, we treated murine splenic naïve CD4+ T cells with and without TMG before activation, polarized towards a Th17 lineage, and then performed an unbiased analysis of the lipidome by mass spectrometry. Of note, cholesterol and total sterol populations were significantly elevated with TMG treatment (Figure 5A). One possibility for these results is increased cholesterol synthesis and thus increased cholesterol biosynthetic intermediates capable of activating RORγt. Additionally, of the top twenty fatty acids detected in the cells, three SFA, such as stearic acid (C18:0), were significantly increased and all PUFA, such as arachidonic acid (C20:4), were significantly decreased with TMG treatment (Figure 5B and 5C). Together, these results demonstrate that elevated O-GlcNAcylation alters the lipid microenvironment such that cholesterol intermediates and fatty acids capable of acting as activating ligands of RORγt at the IL-17A locus were significantly increased. Furthermore, an increase in these lipid ligands is a plausible explanation for why RORγt is retained longer at the IL-17 locus.

**Figure 5.**
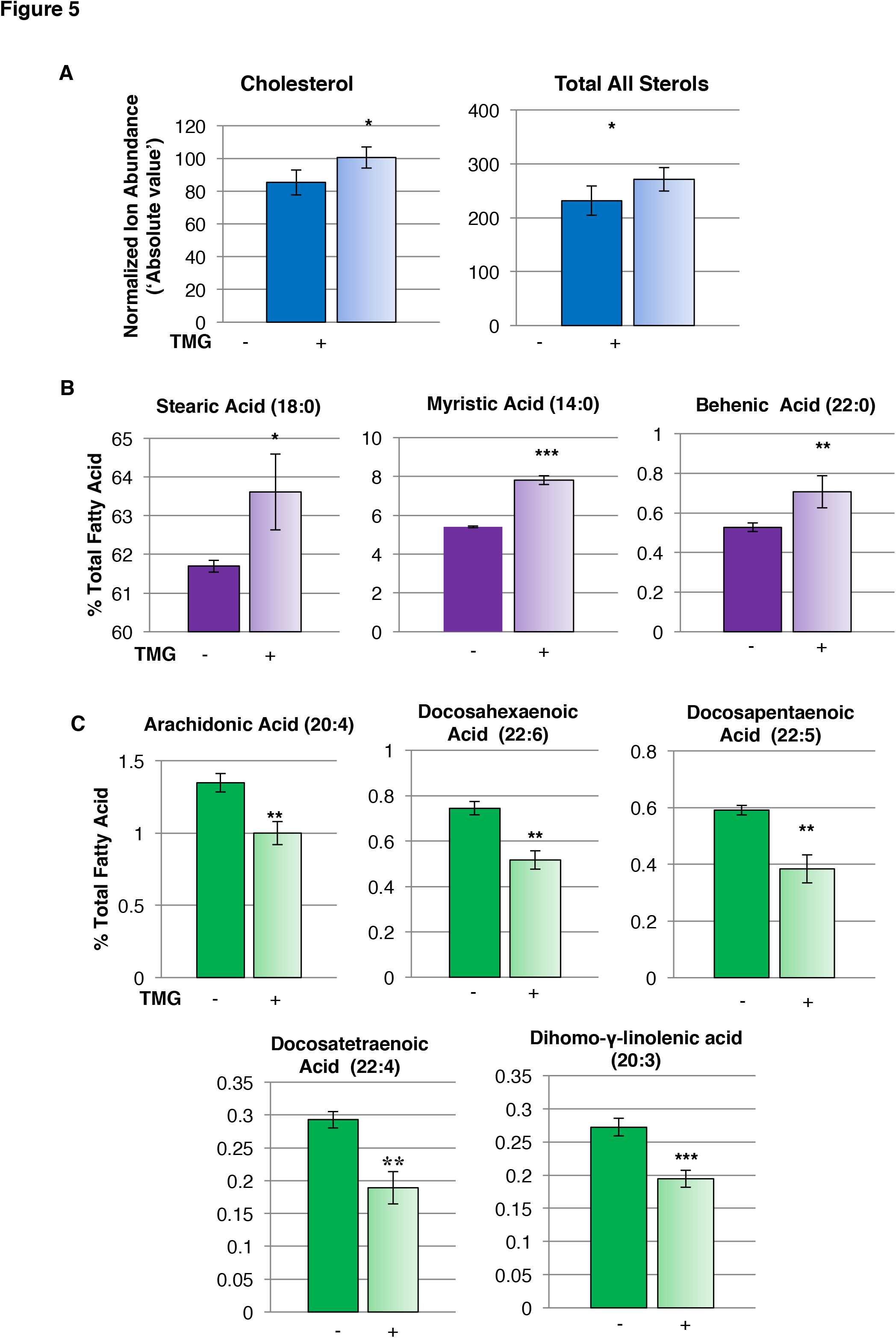
In murine differentiated Th17 cells, elevated O-GlcNAc increases lipid ligands capable of increasing RORγt transcriptional activity. A. The absolute value or normalized ion abundance of total sterols and cholesterol are increased with TMG treatment. B. Of the top 20 fatty acids present in cells, the percentage of saturated fatty acids increased with TMG treatment. C. Of the top 20 fatty acids present in cells, the percentage of polyunsaturated fatty acids decreased with TMG treatment. Bars represent mean +/− SD of 5 biological replicates; * p < 0.05, **p < 0.01, *** p < 0.001

### Acetyl CoA carboxylase 1 (ACC1), the rate limiting enzyme of fatty acid biosynthesis, is modified by O-GlcNAc

The substrates needed for both long chain fatty acids and cholesterol synthesis originate from the activity of acetyl CoA carboxylase (ACC1), the rate-limiting enzyme in fatty acid catabolism. ACC1 catalyzes the carboxylation of acetyl CoA to form malonyl CoA. Importantly, *de novo* fatty acid synthesis is essential for Th17 differentiation (Berod et al., 2014), and ACC1 activity specifically is essential for Th17 differentiation (Endo et al., 2015). Since O-GlcNAcylation and phosphorylation can be reciprocal to one another (Hart et al., 2011), we looked at levels of inhibitory phosphorylation of ACC1 at serine 79 to determine if TMG treatment altered ACC1 inhibition. AMP kinase (5’ adenosine monophosphate-activated protein kinase) senses low ATP levels in a cell and phosphorylates S79 on ACC1 to inhibit biosynthetic activity (Ha et al., 1994). We thus checked phospho-ACC1 levels over 0, 2, 4, 8, 16, and 24 hours after re-stimulation of *ex vivo* differentiated Th17 cells. We found no significant changes in phospho-ACC1 levels (Figure 6A). We next hypothesized that as a cellular nutrient sensor, O-GlcNAc modifies ACC1 and regulates its activity independently of AMPK phosphorylation. We first immunoprecipitated ACC1 from CD4+ T cells and observed that it was O-GlcNAcylated, and the reverse procedure of pulling proteins down with an O-GlcNAc antibody also yielded ACC1 (Figure 6B). We then confirmed this finding by performing electron transfer dissociation mass spectrometry (ETD-MS) on immunoprecipitated ACC1 from TMG treated murine naïve T cells polarized *ex vivo* to the Th17 lineage. Mass spectrometry analysis uncovered 4 high confidence O-GlcNAc sites on murine ACC1: S966, S967, S2091, S2285 (Figure 6C and 6D). Two of these sites (S2091 and S2285) had homologous sites in a reported human partial C-terminal domain ACC1 crystal structure (PDB ID: 4ASI, RCSB Protein Data Bank) (Figure 6D). In the human crystal structure and analogous mouse domains, O-GlcNAc sites were found in the central region (S966 and S976) and carboxyl transferase domain (S2091), implying that O-GlcNAc may regulate binding of protein partners in the central region (Shen and Tong, 2008) and/or the final step in the formation of malonyl CoA respectively, potentially altering ACC1 binding partner function or ACC1 catalytic activity directly.

**Figure 6.**
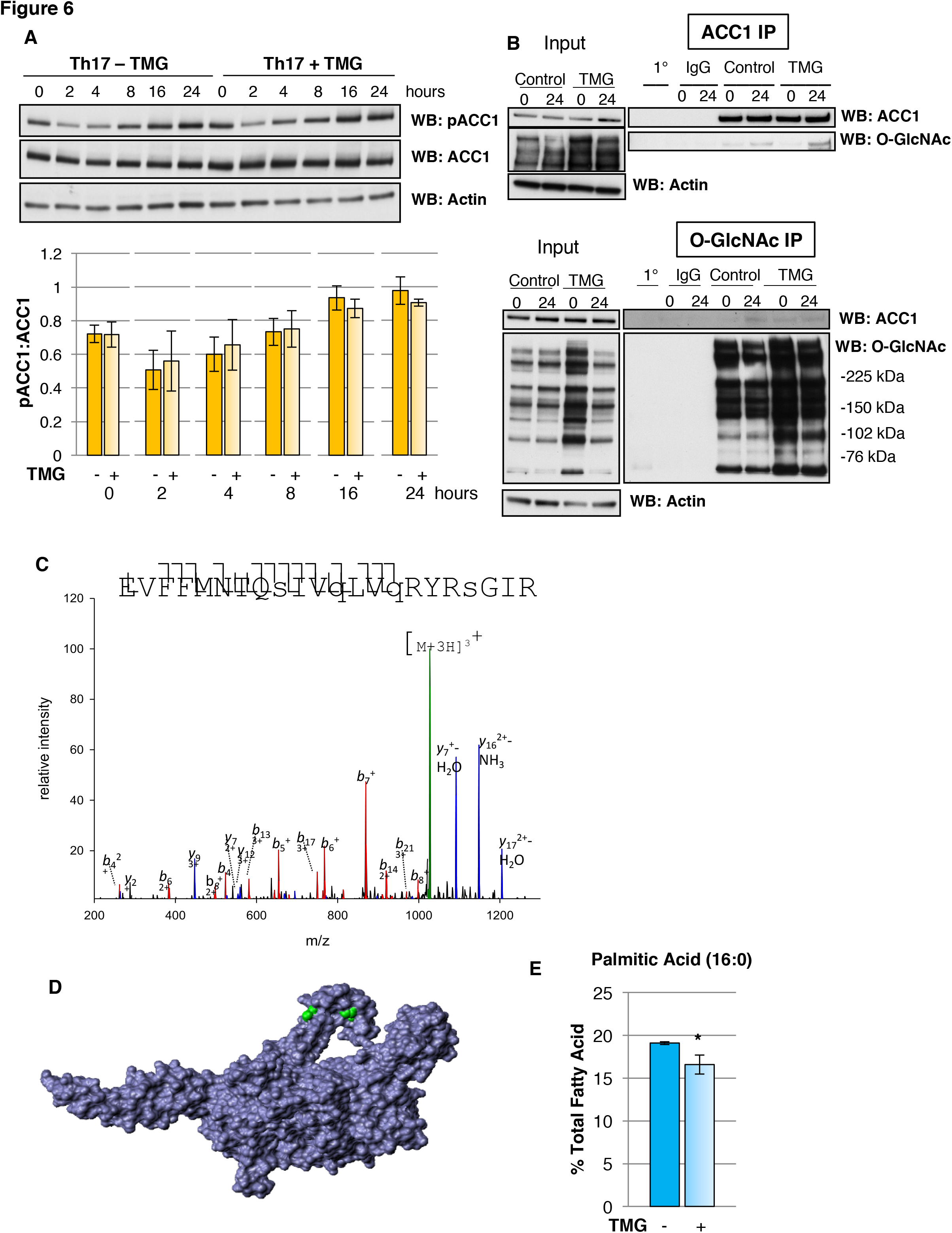
Acetyl CoA carboxylase 1 (ACC1), the rate limiting enzyme of fatty acid biosynthesis, is modified by O-GlcNAc. A. ACC1 or O-GlcNAc modified proteins were immunoprecipitated from CD4+ T cells and show ACC1 is modified by O-GlcNAc. B. Representative mass spectra plot of high confidence O-GlcNAc modified sites S966 and S976 on ACC1. C. Location of analogous O-GlcNAc modified residues S2129 and S2323 in the carboxyl transferase domain on a purified human partial C-terminal crystal structure of ACC1 (PDB #4ASI). D. Inhibitory phosphorylation of ACC1 at S79 is unchanged with TMG treatment over the 24 hours of re-stimulation. pACC1 and ACC1 levels were normalized to actin. Blot is representative of 3 biological replicates. Densitometry bars represent mean +/− SEM of 3 replicates. E. Palmitic acid significantly decreases with TMG treatment. Bars represent mean +/− SD of 5 replicates.

In addition to no change in inhibitory phosphorylation of ACC1, palmitic acid (C16:0), which can negatively feedback to inhibit ACC1 activity, is significantly decreased with OGA inhibition (Figure 6E). Thus, elevated O-GlcNAc levels results in increased lipid ligands that are synthesized downstream of ACC1 activity, and also leads to reduced inhibitory palmitic acid levels and no change in inhibitory phosphorylation of ACC1. Overall, OGA inhibition creates an environment conducive to ACC1 activity and generates lipid ligands capable of increasing RORγt activity at the IL-17 locus.

## DISCUSSION

Obesity has reached epidemic proportions in the developed world and is increasingly becoming a problem in less developed nations (Ng et al., 2014). Patients with diseases of over-nutrition have a well-defined chronic, low-grade inflammation. This inflammation has been implicated in much of the downstream pathology associated with increased adiposity, such as atherosclerosis, insulin resistance, and increased risk of autoimmunity and cancer (Chehimi et al., 2017; Ridker et al., 2017; Schindler et al., 2017). Thus, to lessen the morbidity, mortality, and financial burden consequent to obesity, it is imperative we better understand mechanisms that drive pathology related to excess adiposity. Among this chronic inflammatory milieu, CD4^+^ T cell effectors, particularly Th1 and Th17 cells, are critical players in the development of atherosclerotic plaques and infiltrate adipose tissue to mediate insulin resistance (Chehimi et al., 2017; Winer et al., 2009). In this study, we demonstrate that a homeostatic level of O-GlcNAcylation is necessary for the proper regulation of CD4+ T cell function and differentiation. We cultured murine and human CD4+ T cells with Thiamet-G (TMG), a highly specific O-GlcNAcase inhibitor, to disrupt homeostatic O-GlcNAc levels. We observed a significant increase in pro-inflammatory cytokines, IL-17A and IFNγ, major cytokines secreted by the Th17 and Th1 lineages respectively. We observed no change in IL-10 protein or transcript levels in CD4+ T cells treated with TMG, suggesting no significant decrease in Tregs. Under Th17 polarizing conditions, we observed only a modest, though significant, 4% increase in the number of cells producing IL-17 with TMG treatment. Overall, our data suggests a stronger effect of abnormally elevated O-GlcNAc resulting in more IL-17 being secreted per cell versus skewing differentiation towards Th17s at the expense of Tregs.

Since excess O-GlcNAcylation increased Th17 and Th1 inflammatory cytokines, which are abnormally elevated in obesity, we wanted to determine the role of O-GlcNAc in a Western diet (WD) fed mouse model. In agreement with O-GlcNAc acting as a nutritional sensor, O-GlcNAc levels were significantly increased in the naïve CD4+ T cells of WD fed mice. Interestingly, the levels of OGT and OGA were not different between mice fed standard chow (SC) and mice on the Western diet. Normally, O-GlcNAc levels are vigorously maintained at a level particular for the needs of a cell (Zhang et al., 2014). For example, when O-GlcNAcylation is increased with TMG treatment, OGA levels will increase and OGT levels will decrease to return O-GlcNAc levels to baseline. However, these results suggest that long-term nutritional excess causes O-GlcNAc levels to “reset” to an abnormally high level. When this atypically high new set-point was pushed even further by TMG treatment, both cells from SC and WD fed mice produced significantly more IL-17A. These results are in agreement with a known effect of a single nucleotide polymorphisms (SNPs) in the MGEA gene which encodes OGA. In a Mexican American population with these SNPs, OGA expression is decreased and there is increased incidence and earlier age of onset of type 2 diabetes (Lehman et al., 2005). Alteration to OGT or OGA levels could similarly contribute to an abnormal O-GlcNAc “set point” and pathogenicity. Thus, O-GlcNAcylation is a mechanism contributing to increased pro-inflammatory IL-17A secretion, particularly relevant in the context of diet-induced obesity, where O-GlcNAc levels attain a higher homeostatic level.

Lipid ligands, such as sterols and saturated fatty acids, capable of binding and activating RORγt at the IL-17 locus were increased with TMG treatment. Furthermore, we identified sites of O-GlcNAcylation on acetyl CoA carboxylase 1 (ACC1), which provides the building blocks for both long chain fatty acid and cholesterol synthesis (Kidani and Bensinger, 2017). In addition to increased fatty acid and sterol products that result from ACC1 activity, two other lines of evidence suggest increased ACC1 activity with TMG treatment. First, palmitic acid (C16:0) is the first fatty acid produced as a result of ACC1 initiating fatty acid biosynthesis, and as such, negatively feedbacks to inhibit ACC1. The significant decrease in palmitate with TMG treatment but significant increase in longer chain saturated fatty acids suggests palmitate is being rapidly diverted into elongation pathways or acylation of proteins and is thus less available to inhibit ACC1 activity. Additionally, we were unable to detect a change in inhibitory phosphorylation of ACC1 by AMPK at serine 79, so there is not increased inhibition of ACC1 with TMG treatment. Of note, lipidomics analysis of endothelial cells treated with soraphen A, a potent inhibitor of ACC1, resulted in opposite effects of what we observed (Glatzel et al., 2017). With inhibition of ACC1, polyunsaturated fatty acids (PUFA) increased, whereas we observed a significant decrease in PUFAs in our T cell model. In addition to an O-GlcNAc site (S2091) being present in the carboxyl transferase (CT) domain of ACC1, which catalyzes the final step of malonyl CoA formation, sites were found in the central region of ACC1 (S966 and S976). In a crystal structure of yeast ACC1, the central region was found to be composed of 5 unique domains and was responsible for bringing the biotin carboxylase (BC) and CT domains together (Wei and Tong, 2015), which is an important step in catalysis. Importantly, deletion of residues 940-972 in the first domain of the central region abolished catalytic activity. Overall, multiple lines of evidence suggest TMG treatment favors increased ACC1 activity and production of ligands which enhance RORγt activity.

Obesity and diabetes are major public health concerns and the immune system is a critical driver of pathogenesis in these diseases. Intervening at the level of inflammation is of critical importance in preventing pathological complications. Unfortunately, the mechanisms driving chronic inflammation have been difficult to elucidate. The results of this study represent the first investigation of how the O-GlcNAc post-translational modification regulates CD4+ T cell function and differentiation and support a role for aberrant O-GlcNAcylation in mediating a pro-inflammatory Th17 response in a diet-induced obesity mouse model. Human CD4+ T cells showed a similar significant increase in IL-17 production when treated with TMG. Thus, mechanistic insights derived from murine CD4+ T cells may be applicable to humans. Importantly, we discovered that the rate-limiting enzyme in fatty acid biosynthesis, ACC1, is O-GlcNAcylated and the lipid profile of cells treated with TMG shifts towards an inflammatory phenotype and produces ligands driving RORγt transcriptional activity at the IL-17 locus. This works thus fills in a piece of the mechanistic puzzle for why pathogenic inflammatory cytokine production increases in CD4+ T cells in diseases of over-nutrition.

## ACKNOWLEDGEMENTS

This work was supported by National Institute of Diabetes and Digestive and Kidney Diseases grant R01DK091277 awarded to P. Fields, National Institute of Diabetes and Digestive and Kidney Diseases grant R01DK100595 awarded to C. Slawson, the Molecular Regulation of Cell Development and Differentiation COBRE P30GM122731 awarded to P. Fields and C. Slawson, and a KUMC Biomedical Research Training Program grant awarded to M. Machacek. We acknowledge the Flow Cytometry Core Laboratory, which is sponsored, in part, by the NIH/NIGMS COBRE grant P30 GM103326 and especially thank Dr. Rich Hastings, Flow Core Director, for his invaluable assistance and expertise. We acknowledge support from the University of Kansas (KU) Cancer Center’s Biospecimen Repository Core Facility staff for helping obtain human specimens, which receives support from the KU Cancer Center’s Cancer Center Support Grant (P30 CA168524). Finally, we are indebted to the volunteers who generously donated blood in support of this study.

## AUTHOR CONTRIBUTIONS

Conceptualization, P.F., C.S., and M.M.; Methodology, P.F. C.S., T.Li, and M.M.; Investigation, M.M., Z.Z., E.T., J.L., M.V., A.A., and T. Lydic; Writing –Original Draft, M.M‥; Writing –Review & Editing, all authors; Funding Acquisition, P.F. and C.S.; Resources, J.L., T.Li; Supervision, P.F. and C.S.

## MATERIALS & METHODS

### Mouse T cell isolation, activation, and differentiation

C57BL/6 male (Western diet experiments) and female mice (Jackson Laboratories) between 6-24 weeks of age were used in these studies. Mice were humanely euthanized using CO_2_ asphyxiation according to Institutional Animal Care and Use Committee (IACUC) approved protocol. CD4+ T cells were isolated from spleen using the CD4+ T cell isolation kit (Stem Cell Technologies). CD4+ T cells alone or naïve cells obtained by staining with APC-CD62L and FITC-CD44 (BD Biosciences) were sorted and treated with 10 μM Thiamet-G (SD Chemmolecules, LLC) or vehicle control for 6 hours. Cells (2.5 × 10^5^ cells/well) were then activated on anti-CD3 and anti-CD28 (Bio X Cell) antibody coated plates. Plates were prepared by addition of 2 μg anti-CD3 and 1 μg anti-CD28 per well of a 48-well plate in PBS containing calcium and magnesium incubated overnight at 4°C. At the time of activation, polarizing cytokines were added to cells to promote differentiation towards specific lineages as follows: Th0 - no cytokines added; Th17 - mouse IL-6 (20 ng/mL, Miltenyi), human TGFβ1 (2 ng/mL, Miltenyi), anti-IL-4 (10 μg/mL), anti-IFNγ (10 μg/mL). All cells were incubated at 37 °C in 5% CO_2_ in a 95% humidified incubator. On the second day of culture, mouse IL-2 (eBioscience) at 20 ng/mL was added to cultures and the cells were re-suspended. On the fourth day of culture, cells were re-suspended, any dead cells were removed by gradient centrifugation with Lymphocyte Separation Medium (Corning), and counted with a hemocytometer. 2.5 × 10^5^ cells per well were then re-stimulated on plates containing 3 μg anti-CD3 and 1 μg anti-CD28 per well, which were prepared as previously described. 24 hours after re-stimulation supernatants and cells were harvested. Mouse IL-17A, IFNγ, IL-4, or IL-10 levels in the supernatants were quantified using the cytokine’s corresponding ELISA kit (eBioscience).

### Western Diet Mouse Model

For Western diet studies, 6-8 week old male C57BL/6 mice (Jackson Laboratories) were fed a 42% kcal from fat (>60% saturated fat), 0.2% cholesterol, and high sucrose “Western diet” (Teklad TD.88137) for 16 weeks. Control 6-8 week old male C57BL/6 mice were fed standard chow, containing 14% kcal from fat (Teklad 8604). Mice were weighed at 0, 2, 4, 6, 8, 10, 12, 14, and 16 weeks. At 15 weeks, control and Western diet fed mice were fasted overnight for 15 hours. At the end of the fast, tail vein blood was used to determine the blood glucose level (mg/dL) using One Touch Ultra Blue test strips and the One Touch UltraMini blood glucose monitoring system (LifeScan). At 16 weeks mice were sacrificed following a 15 hour overnight fast, and splenic CD4+ T cells were used for subsequent studies as described.

### Intracellular Cytokine Staining

CD4+ T cells were isolated and cultured as previously described. After four days of culture, cells were re-suspended, any dead cells were removed by gradient centrifugation with Lymphocyte Separation Medium (Corning), and viable cells were re-suspended at a concentration of 1-10 million cells per mL. Cells were stimulated with 50 ng/mL PMA (phorbol 12-myristate 13-acetate) and 1 μg/mL ionomycin (both Sigma) for 5 hours. Protein secretion was inhibited by addition of 4 μL of BD GolgiStop™ Protein Transport Inhibitor (containing monensin) per 6 mL of culture. After stimulation, cells were washed twice with PBS + 2% FBS and counted. One million cells per condition were added to an Eppendorf tube, spun, and re-suspended in cold BD Cytofíx™ buffer from the Mouse Th1/Th17 Phenotyping Kit (BD Biosciences) and incubated for 20 minutes at room temperature. Cells were then washed twice with PBS + 2% FBS and diluted in BD Perm/Wash Buffer™ for 15 minutes at room temperature. Cells were centrifuged and then re-suspended in 50 μL BD Perm/Wash Buffer™ plus 20 μL/tube of staining cocktail (mouse PerCP-Cy5.5-CD4, PE-IL-17A, FITC-IFNγ) or BD Perm/Wash Buffer™ as a negative control or mouse PerCP-Cy5.5-CD4 and isotype controls for IL-17A and IFNγ for 30 minutes at room temperature. Cells were washed twice with PBS + 2% FBS and data was acquired on an LSR II (BD Biosciences) and analyzed with FlowJo (Tree Star).

### Quantitative Real Time Polymerase Chain Reaction (qPCR)

Mouse or human RNA was extracted by dissolving 2 × 10^6^ cells in TRI reagent (Sigma) followed by the addition of chloroform. The mixture was mixed by inversion, centrifuged, and the aqueous layer was taken. Isopropanol was added to the aqueous layer to precipitate the RNA. The RNA was pelleted by centrifugation, washed with 75% ethanol, and allowed to air-dry before adding DEPC-treated, RNase free water (Ambion). RNA of high purity (260/280 and 260/230 greater than 1.7) was used for further analysis. 0.5-0.75 μg RNA was converted to cDNA using iScript reverse transcriptase reaction mix (Biorad) in a thermal cycler (Model 2720, Applied Biosystems) using the following protocol: priming, 5 min at 25 °C; RT, 30 min at 42 °C; and RT inactivation, 5 min at 85 °C. For qPCR reactions, 10 μL SsoAdvanced Universal SYBR Green Supermix (Biorad), 10-100 ng of cDNA, 0.2 μL each of forward and reverse primers (100 μM stock concentration), and water to a final reaction volume of 20 μL were added per well to 96 well-plates (Midsci). The reactions were run on a CFX96 Real Time PCR Detection System (Biorad), using the following conditions: polymerase activation and DNA denaturation, 30 s at 95 °C; amplification denaturation, 5 s at 95 °C; and amplification annealing and extension, 30 s at 60 °C or 62 °C for 40 cycles. The dynamic range of reverse transcription and the amplification efficiency was determined for each primer pair and cell culture condition. Thus, Cq values within these ranges were reliably used to calculate the fold change in gene expression compared to an internal standard using the ^ΔΔ^Cq method.

### Immunoblotting

Cells were lysed on ice in 20 mM Tris, pH 7.4, 150 mM NaCl, 40 mM GlcNAc, 2 mM EDTA, 1 mM DTT, 1% Nonidet P-40 lysis buffer with protease inhibitors (1 mM β-glycerophosphate, 1 mM NaF, 2 mM PMSF, 1X inhibitor cocktail composed of 1 μg/ml leupeptin, 1 μg/ml antipain, 10 μg/ml benzamidine, and 0.1% aprotinin (Sigma)) added immediately before lysis. Lysates were incubated on ice for 20 minutes and vortexed every 5 minutes. Lysates were then centrifuged for 20 minutes, 4°C and the supernatant was removed to another tube. Protein concentration of the lysate was determined using protein assay dye reagent (Biorad), using known concentrations of bovine serum albumin (Biorad) as the standard. Lysates were then denatured by addition of 4X protein solubility mixture (100 mM Tris, pH 6.8, 10 mM EDTA, 8% SDS, 50% sucrose, 5% beta-mercaptoethanol, 0.08% Pyronin Y) and boiling for 2 minutes. Equal amounts of lysates were loaded onto 4-15% Criterion precast TGX gels (Biorad). Electrophoresis occurred at 125V, and then the gel proteins were transferred to PVDF membranes at 0.4A. Membranes were blocked with 3% BSA, 0.01% sodium azide in TBST (25 mM Tris, pH 7.6, 150 mM NaCl, 0.05% Tween-20) for at least 20 minutes. Blots were then probed overnight at 4°C with primary antibody to protein of interest at 1:1,000 dilution. The next day blots were washed 5 times in TBST for 5 minutes each. HRP-conjugated secondary antibody at 1:10,000 dilution was added for an hour at room temperature, followed by washing 5 times for 5 minutes each. Blots were then developed using the chemiluminescence HRP (horseradish peroxidase) antibody detection method (HyGlo Spray, Denville Scientific). Commonly, blots were striped with 200 mM glycine, pH 2.5 for one hour at room temperature, blocked, and re-probed with antibodies to other proteins of interest. Where shown, Image J (Schneider et al., 2012) was used to quantify the density of protein bands compared to an internal standard protein band such as actin. Antibodies used in these studies: pACC1 (Ser79), ACC1, pSTAT3 (Tyr705), STAT3 (Cell Signaling Technologies); O-GlcNAc (clone: RL2), RORγt (Abcam); actin (Sigma); OGT (clone: AL-34) and OGA (clone: 345) antibodies were a generous gift from the laboratory of Dr. Gerald Hart in the Department of Biological Chemistry at The Johns Hopkins University.

### Immunoprecipitation (IP)

CD4+ T cells were lysed with lysis buffer (50 mM Tris, pH 7.4, 150 mM NaCl, 1 mM DTT, 1% NP-40, 40 mM N-acetylglucosamine) and protein concentration was quantified using Protein Assay Dye Reagent (Biorad). One μg of antibody for the protein of interest or one μg of isotype matched IgG was added to equal amounts of whole cell lysate diluted to an equal volume in Nonidet P-40 lysis buffer and rotated overnight at 4°C. As an additional control, 1 μg antibody to the protein of interest was added to lysis buffer only and rotated overnight at 4°C. Protein G-Sepharose beads (Millipore) were added and rotated at 4°C for 2 hours. The protein G beads were then washed three times with NP-40 lysis buffer containing 600 mM NaCl followed by two PBS washes, briefly vortexing beads in between washes. Washed beads were re-suspended in Laemlli buffer, boiled for 2 minutes, and run on 4-15% gradient Criterion TGX gels (Biorad). Gel proteins were transferred to PVDF membranes and blocked with 3% BSA, 0.01% sodium azide in TBST (25 mM Tris, pH 7.6, 150 mM NaCl, 1% Tween-20). Blots were then probed overnight at 4°C with antibodies to O-GlcNAc (RL2, Abcam) followed by immunopreciptated protein (ACC1 (CST) and RORγt (Abcam)) and developed using the chemiluminescence HRP (horseradish peroxidase) antibody detection method (HyGlo Spray, Denville Scientific). For RORγt immunoprecipitation, samples were treated with 5 units of IdeZ protease (Promega) in PBS for 30 minutes at 37°C after washing the beads.

### Chromatin Immunoprecipitation (ChIP)

Mouse naïve CD4+ T cells were isolated as previously described from mice fed standard chow. TMG was added to naïve CD4+ T cells for 6 hours before activation and differentiated towards the Th17 lineage as previously described. After 4 days of culture, cells were harvested and fixed with 2 mM ethylene glycol bis(succinimidyl succinate) (EGS; Thermo Scientific) in PBS for 30 minutes followed by 1% formaldehyde (Thermo Scientific) for 10 minutes. Nuclei were extracted by re-suspending fixed cells in hypotonic cell lysis buffer (10 mM Tris, pH 8.0, 10 mM NaCl, 3 mM MgCl2, 40 mM GlcNAc, 0.5% NP-40) with freshly added protease inhibitors (1 mM β-glycerophoshate, 1 mM NaF, 1X protease inhibitor cocktail (containing aprotinin, leupeptin, antipain, benzamidine), and 2 mM PMSF) for 20 minutes on ice. Nuclear lysis buffer (50 mM Tris, pH 8.0, 10 mM EDTA, 1% SDS, 25% glycerol) with freshly added protease inhibitors as previously described and the chromatin was sonicated using the QSonica sonicator and cooling system (Model #:Q800R and Oasis 180 respectively) for 200 cycles of 15 seconds on at 75% amplitude and 45 seconds off. Sonication efficiency was checked by treating sheared chromatin diluted in ChIP buffer (50 mM Tris, pH 7.5, 150 mM NaCl, 5 mM EDTA, 0.5% NP-40, 1% Triton X-100) with 1 μg RNase A (Invitrogen) overnight at 65°C, followed by 2 ug Proteinase K (Sigma) for 6 hours at 65°C, before addition of phenol:chloroform:isoamyl alcohol (Acros Organics). The top aqueous layer was taken and 20 μg of glycogen (Invitrogen) followed by 1 mL of cold 100% ethanol (Sigma) was added and incubated overnight at −20°C to precipitate the DNA. The DNA was pelleted by centrifuging for 30 minutes at 4°C, then washed with 75% ethanol, and centrifuged for an additional 10 minutes at 4°C. The supernatant was aspirated and the DNA pellet was allowed to dry for at least 6 hours. DNA was re-suspended in DEPC-treated, DNase free water (Ambion) and run on a 1.5% agarose gel. Bands of 100-300 base pairs in length indicated a successful sonication cycle. After ensuring adequate shearing of the chromatin, 2 μg of antibody to the protein of interest or 2 ug of isotype control was then added to equal amounts of sheared chromatin diluted four-fold with ChIP buffer and rotated overnight at 4°C. The next day 20 μL of isotype matched Dynabeads (Life Technologies) were added and rotated for another 2 hours. Beads were then washed with 1 mL cold ChIP buffer five times using the DynaMag-2 Magnet (12321D, Invitrogen), rotating for 5 minutes at 4°C between washes. 100 μL of 10% Chelex 100 slurry was added to the beads and boiled for 10 minutes. 1 μg of RNase A was added and incubated at 65°C for 15 minutes minimum followed by addition of 2 μg Proteinase K for 30 minutes at 65°C. The samples were boiled again for 10 minutes to inactivate Proteinase K, and then centrifuged at 12,000 × g for one minute, 4°C. The supernatant (70 μL) was transferred to a new tube and the remaining Chelex resin was washed with 130 μL water, re-spun, and resulting supernatant added to the first one. 10% of the chromatin used for the IP was reserved as input for the qPCR reaction. DNA from inputs was extracted using the same method to determine sonication efficiency. 10 μL of the immunoprecipitated DNA or DNA purified from inputs was used per reaction for quantitative PCR and run on the CFX96 Real Time PCR Detection System as previously described. The Cq values were normalized to percentage of input. Antibodies used in these studies: rabbit control IgG and RORγt (Abcam)

### Lipidomics

Mouse naïve CD4+ T cells were isolated, activated, and differentiated towards the Th17 lineage as previously described from mice fed standard chow with and without TMG treatment. After four days of culture, cells were re-stimulated with plate-bound anti-CD3 and anti-CD28 for 8 hours before harvesting, washing with PBS, and snap-freezing the cell pellet.

#### Lipid extraction.

Frozen cell pellets were subjected to monophasic lipid extraction in methanol: chloroform: water (2: 1: 0.74, v: v: v) as previously described (Lydic et al., 2014). During lipid extraction, each sample was spiked with synthetic di-myristoyl phosphatidylcholine obtained from Avanti Polar Lipids (Alabaster, AL), and D7-cholesterol obtained from Steraloids (Newport, RI) at 1 nmole/mg protein as an internal standard for relative quantitation of lipids and sterols. Dried lipid extracts were washed three times with 10 mM ammonium bicarbonate, dried under vacuum, and resuspended in methanol containing 0.01% butylated hydroxytoluene. Samples were stored under a blanket of nitrogen at −80 degrees Celsius until further analysis.

#### Global Lipidomics Analysis

For each analysis, lipid extracts were transferred to an Eppendorf twin-tec 96-well plate (Sigma Aldrich, St. Louis, MO), and evaporated under nitrogen. The dried lipid film was then resuspended in isopropanol:methanol:chloroform (4:2:1 v:v:v) containing 20 mM ammonium formate and sealed with Teflon Ultra Thin Sealing Tape (Analytical Sales and Services, Pompton Plains, NJ). Appropriate sample dilution to minimize ion suppression effects was determined as previously described (Lydic et al., 2014). Global lipidomics analysis was performed by direct infusion high resolution/accurate mass spectrometry and tandem mass spectrometry. Samples were introduced to the mass spectrometer by nanoelectrospray ionization (nESI) using an Advion Triversa Nanomate nESI source (Advion, Ithaca, NY) with a spray voltage of 1.4 kV and a gas pressure of 0.3 psi. A Thermo Scientific LTQ-Orbitrap Velos (San Jose, CA) mass spectrometer was used for lipid detection. High resolution MS and MS/MS spectra were acquired for each sample in positive and negative ionization modes using the Orbitrap FT analyzer operating at 100,000 mass resolving power (defined at m/z 400).

#### Analysis of Free and Total Sterol Content

Sterols and oxysterols were analyzed by high resolution/accurate mass LC-MS using a Shimadzu Prominence HPLC equipped with an in-line solvent degassing unit, autosampler, column oven, and two LC-20AD pumps, coupled to a Thermo Scientific LTQ-Orbitrap Velos mass spectrometer. Lipid extracts were used directly for analysis of ‘free’ sterols, or subjected to alkaline hydrolysis of sterol esters for analysis of total cellular sterols as previously described (McDonald et al., 2007). Ten microliter injections of each total or hydrolyzed lipid extract were separated using a Phenomenex (Torrance, CA) Synergi Hydro-RP C18 column, 2 mm × 150mm, with 3 micron particles and 80 micron pore size, at 50 degrees Celsius. The gradient elution method was modified from (McDonald et al., 2007) and utilized A: water containing 0.1% formic acid and B: methanol containing 0.1% formic acid, with a 13 minute gradient from 85% B to 100% B at a flow rate of 250 microliters/minute. The column eluent was introduced to the LTQ-Orbitrap Velos by a heated electrospray ionization source, using a spray voltage of 4.5 kV, sheath gas pressure of 30 psi, and auxillary gas pressure of 10 psi. The source heater and MS capillary inlet tube were held at 350 degrees Celsius. Sterols and oxysterols were analyzed by full scan positive ion mode MS using the Orbitrap detector at a resolution of 60,000 (defined at m/z 400), to detect sterol in-source fragmentation derived [M-H_2_0+H]^+^ and [M-2H_2_0+H]^+^ ions, which were the most abundant sterol ions formed under the conditions utilized. Targeted high resolution HCD-MS/MS at 45% normalized collision energy was employed on known sterol-related ions and their product ions (McDonald et al., 2007) to verify the identity of each sterol peak.

#### Peak finding, analyte identification, and quantification

For global lipidomics analysis, lipids were identified using the Lipid Mass Spectrum Analysis (LIMSA) v.1.0 software linear fit algorithm, in conjunction with a user-defined database of hypothetical lipid compounds for automated peak finding and correction of ^13^C isotope effects. Relative quantification of lipid abundance between samples was performed by normalization of target lipid ion peak areas to the di-myristoyl phosphatidylcholine internal standard as previously described (Lydic et al., 2015). For free and total sterol analysis, chromatographic peak alignment, compound identification, and relative quantitation against the D7-cholesterol internal standard was performed using MAVEN software (Clasquin et al., 2002).

### Mass spectrometry

Mouse naïve CD4+ T cells were isolated, activated, and differentiated towards the Th17 lineage as previously described from mice fed standard chow. After 4 days of culture, cells were harvested, lysed, and ACC1 was immunoprecipitated as previously described. Immune-purified complexes bound to agarose beads were trypsin-digested as previously described (Mohammed et al., 2013).

#### LC-MS analysis.

Following enzymatic digestion, the resulting peptides were concentrated on a centrivac concentrator (Labconco) to a final volume of 20 μL. The peptide extracts were analyzed using an Orbitrap Fusion Lumos mass spectrometer (Thermo Fisher Scientific) coupled to an uHPLC (nLC 1200, Thermo Fisher Scientific). In each run, the sample was injected directly into the reversed phase column (Acclaim PepMap RSLC, 50 μm × 15 cm, C18 reversed phase 2 μm, 100Å), and washed with 0.1% formic acid for 5 minutes at 500 nl/min. The column was mounted on the electrospray stage of the mass spectrometer, and the peptides were eluted with a gradient of acetonitrile (solvent B: 90 % acetonitrile, 0.1% formic acid) in 0.1% formic acid in water (solvent A). The gradient profile was as follows: 0-5 min, 5-15% B; 5-60 min, 15-40% B; 60-70 min, 40% B; 70-75 min, 40-75% B; 75-83 min 75% B. At the end of each chromatographic run, the column was washed with 100% B. The flow rate was maintained at 300 nl/min.

#### MS Acquisition.

The mass spectrometer was operated in positive ionization mode with a nanospray flex ionization source operated at 2.6 Kv and source temperature at 250. Multiple runs were performed using different fragmentation methods as described below. The instrument was operated in data-dependent acquisition mode, with full MS scans over a mass range of m/z 350-1900 with detection in the Orbitrap (120 K resolution) and with auto gain control (AGC) set to 100,000. In each cycle of data-dependent acquisition analysis, following each survey scan, the most intense ions above a threshold ion count of 30,000 were selected for fragmentation at normalized collision energy of 28% (HCD) or 35% (CID). The number of selected precursor ions for fragmentation was determined by the “Top Speed” acquisition algorithm and a dynamic exclusion of 60 s. Fragment ion spectra were acquired in the linear IT, with an AGC of 4,000 and a maximum injection time of 300 ms for IT MS2 detection. For ETD fragmentation, calibrated charge dependent ETD parameters were enabled to determine ETD reagent ion AGC and ETD reaction times, and all MS/MS were mass analyzed in the Orbitrap with a resolution of 15,000 K at 200 m/z. All data were acquired with Xcalibur software v3.0.63 (Tune v2.0 1258).

#### Data analysis

For data analysis all MS/MS scans were searched using Proteome Discoverer (version 2.2, Thermo Fisher Scientific) running the Sequest HT algorithm. A database search was conducted against a human protein database derived from the NIBInr repository as in January 9, 2017. The refined data were subjected to database search using trypsin cleavage specificity, with a maximun of 2 missed cleavages. The following variable modifications were selected: pyroglutamination from Q and E (N-terminal), oxidation of M, and deamidation of N, Q. Carboxymethylation of C was selected as a fixed modification. HexNac was selected as variable modification. A maximum of 3 modifications/peptide was allowed. Estimation of false positive rate (FDR) was conducted by searching all spectra against a decoy database consisting of the inverted sequences of all proteins in the original (direct) database. For peptide identification a FDR £ 1 was defined and a minimum of two unique peptides per protein was required for protein identification. Amino acid sequence assignment of all peptides of interest was subsequently inspected manually.

### Human T cell assays

Human whole blood samples were obtained with consent from patient volunteers by the Biospecimen Repository Core Facility (BRCF) at the University of Kansas Medical Center. Inclusion criteria for volunteers was a body mass index (BMI, calculated as weight in kilograms/height in meters squared) between 18-30 and no prior type 1 or type 2 diabetes diagnoses. Exclusion criteria for samples included patient history of or current immunotherapy for cancer, autoimmune disorder, or current use of steroids or immune-modifying drugs (i.e. methotrexate). Whole blood was diluted 1:1 with PBS and gently layered on top of Lymphocyte Separation Medium (Corning) and then centrifuged at 2,000 × g for 15 minutes at room temperature with the lowest level of brake. Samples of plasma were collected and snap frozen, and the remaining plasma was aspirated. The white blood cell layer was collected and washed twice with PBS. CD4+ T cells were isolated from the washed white blood cell layer using antihuman CD4 particles (BD Biosciences). Naïve CD4+ T cells were obtained by staining CD4+ cells with mouse anti-human CD45RA and mouse anti-human CD197 (CCR7) (BD Biosciences) for 30 minutes on ice, washing with PBS + 2% FBS, and then sorting for the double-positive population of cells. Unsorted CD4+ T cells and naïve cells were treated with TMG or vehicle for 6 hours before activation with anti-CD3 and anti-CD28 conjugated beads from the Human T Cell Activation/Expansion Kit (Miltenyi). Cells were cultured in RPMI-1640 media (Sigma) containing 10% FBS (Gemini), 1% penicillin-streptomycin (Life Technologies), 2 mM GlutaMAX (Life Technologies), 10 mM HEPES, 1 mM sodium pyruvate, and 1 mM MEM nonessential amino acids (Sigma). Unsorted CD4+ T cells and naïve CD4+ T cells were cultured under both Th0 (no polarizing cytokines added) or Th17 conditions (human IL-6 (50 ng/mL), human IL-1β (10 ng/mL), human IL-23 (10 ng/mL), human TGFβ1 (3 ng/mL) (all Miltenyi), and neutralizing antibodies to IFNγ (clone: XMG1.2) and IL-4 (clone: 11B11) at 10 μg/ml (Bio X Cell)). All cells were incubated at 37 °C in 5% CO2 in a 95% humidified incubator. On day 3 of culture, 20 ng/mL of human IL-2 (Miltenyi) was added to cells. On day 7 of culture, cells and supernatants were harvested. IL-17A levels in supernatants were quantified using a human IL-17A ELISA (Invitrogen) and normalized to number of cells present at day 7. Protein and RNA was harvested from cells and used for western blotting or qPCR analyses respectively as previously described.

### Statistics

Student’s T test was performed to compare means between two groups. A p value less than 0.05 was considered statistically significant (*), less than 0.01 (**), and less than 0.001 (***). Standard deviation or standard error of the mean is used for error bars as indicated in figure legends.

## SUPPLEMENTAL INFORMATION

Primer Tables

**Figure S1.**
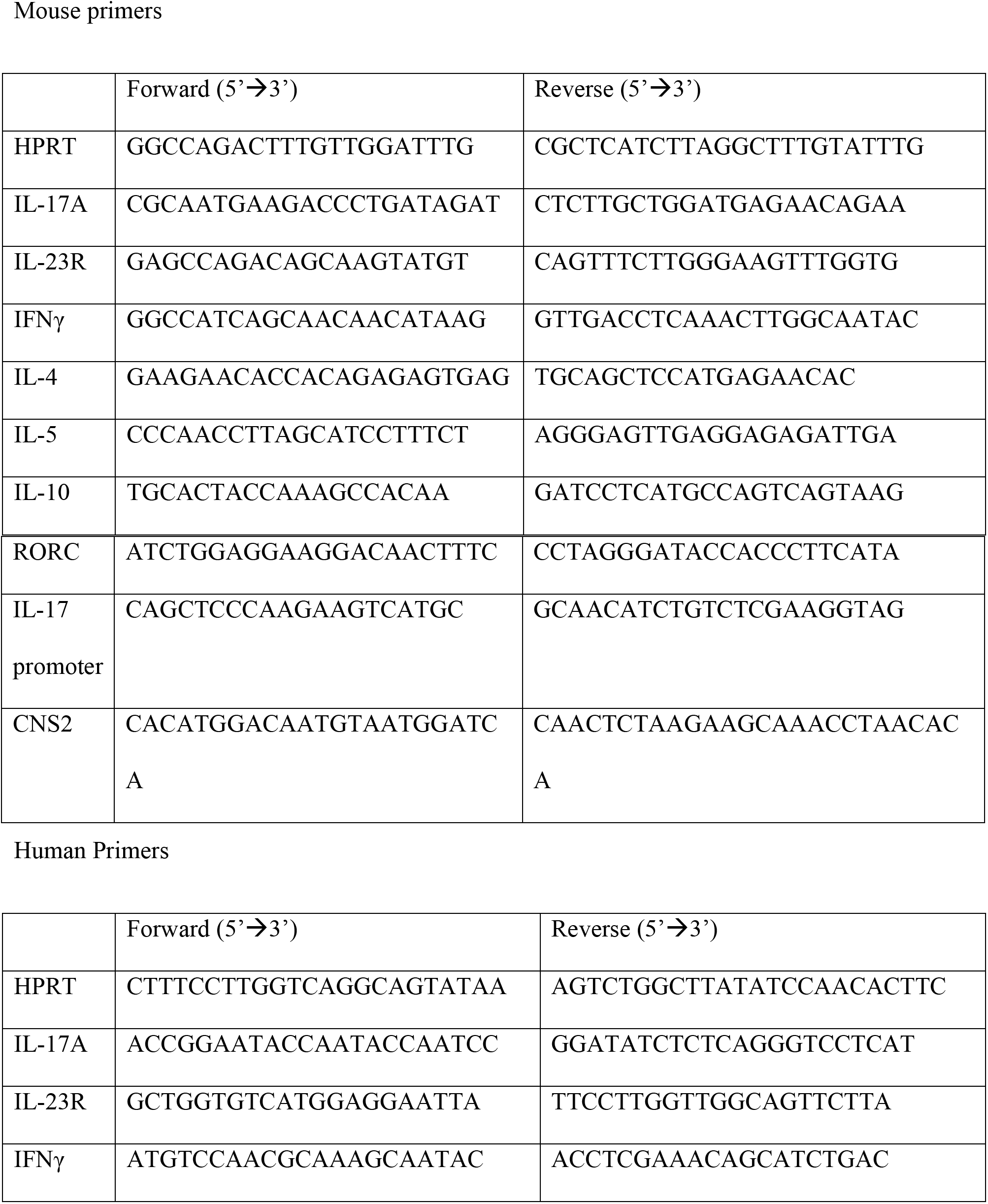
Th17 cells differentiated *ex vivo* from standard chow and Western diet fed mice have no change in protein or transcript levels of RORγt. A. After restimulation for 24 hours on the fourth day of culture, protein levels of RORγt are unchanged. B. After restimulation for 24 hours on the fourth day of culture, RORγt transcript levels are unchanged.

## REFERENCES

Aggarwal, S., Ghilardi, N., Xie, M.-H., de Sauvage, F.J., and Gurney, A.L. (2003). Interleukin-23 Promotes a Distinct CD4 T Cell Activation State Characterized by the Production of Interleukin-17. Journal of Biological Chemistry 278, 1910–1914.

Berod, L., Friedrich, C., Nandan, A., Freitag, J., Hagemann, S., Harmrolfs, K., Sandouk, A., Hesse, C., Castro, C.N., Bahre, H.,et al. (2014). De novo fatty acid synthesis controls the fate between regulatory T and T helper 17 cells. Nat Med 20, 1327–1333.

Chang, C.-H., Curtis, Jonathan D., Maggi, Leonard B., Jr., Faubert, B., Villarino, Alejandro V., O’Sullivan, D., Huang, Stanley C.-C., van der Windt, Gerritje J.W., Blagih, J., Qiu, J.,et al. (2013). Posttranscriptional Control of T Cell Effector Function by Aerobic Glycolysis. Cell 153, 1239–1251.

Chehimi, M., Vidal, H., and Eljaafari, A. (2017). Pathogenic Role of IL-17-Producing Immune Cells in Obesity, and Related Inflammatory Diseases. Journal of Clinical Medicine 6, 68.

Chen, Z., Tato, C.M., Muul, L., Laurence, A., and O’Shea, J.J. (2007). Distinct regulation of interleukin-17 in human T helper lymphocytes. Arthritis & Rheumatism 56, 2936–2946.

Clasquin, M.F., Melamud, E., and Rabinowitz, J.D. (2002). LC-MS Data Processing with MAVEN: A Metabolomic Analysis and Visualization Engine. In Current Protocols in Bioinformatics (John Wiley & Sons, Inc.).

Eljaafari, A., Robert, M., Chehimi, M., Chanon, S., Durand, C., Vial, G., Bendridi, N., Madec, A.-M., Disse, E., Laville, M.,et al. (2015). Adipose Tissue-Derived Stem Cells From Obese Subjects Contribute to Inflammation and Reduced Insulin Response in Adipocytes Through Differential Regulation of the Th1/Th17 Balance and Monocyte Activation. Diabetes 64, 2477–2488.

Endo, Y., Asou, Hikari K., Matsugae, N., Hirahara, K., Shinoda, K., Tumes, Damon J., Tokuyama, H., Yokote, K., and Nakayama, T. (2015). Obesity Drives Th17 Cell Differentiation by Inducing the Lipid Metabolic Kinase, ACC1. Cell Reports 12, 1042–1055.

Frauwirth, K.A., Riley, J.L., Harris, M.H., Parry, R.V., Rathmell, J.C., Plas, D.R., Elstrom, R.L., June, C.H., and Thompson, C.B. (2002). The CD28 Signaling Pathway Regulates Glucose Metabolism. Immunity 16, 769–777.

Geginat, J., Paroni, M., Facciotti, F., Gruarin, P., Kastirr, I., Caprioli, F., Pagani, M., and Abrignani, S. (2013). The CD4-centered universe of human T cell subsets. Seminars in Immunology 25, 252–262.

Ghoreschi, K., Laurence, A., Yang, X.-P., Tato, C.M., McGeachy, M.J., Konkel, J.E., Ramos, H.L., Wei, L., Davidson, T.S., Bouladoux, N.,et al. (2010). Generation of pathogenic TH17 cells in the absence of TGF-[bgr] signalling. Nature 467, 967–971.

Glatzel, D.K., Koeberle, A., Pein, H., Loeser, K., Stark, A., Keksel, N., Werz, O., Mueller, R., Bischoff, I., and Fuerst, R. (2017). Acetyl-CoA carboxylase 1 regulates endothelial cell migration by shifting the phospholipid composition. Journal of Lipid Research.

Golks, A., Thi-Thanh Thao Tran, Jean Francois Goetschy, and Danilo Guerini (2007). Requirement for O-linked N-acetylglucosaminyltransferase in lymphocytes activation. European Molecular Biology Organization 26, 4368–4379.

Golks, A., Tran, T.T.T., Goetschy, J.F., and Guerini, D. (2007). Requirement for O-linked N-acetylglucosaminyltransferase in lymphocytes activation. The EMBO Journal 26, 4368–4379.

Ha, J., Daniel, S., Broyles, S.S., and Kim, K.H. (1994). Critical phosphorylation sites for acetyl-CoA carboxylase activity. Journal of Biological Chemistry 269, 22162–22168.

Hart, G.W., and Kelly P. Kearse (1991). Topology of O-linked N-acetylglucosomaine in murine lymphocytes. Archives of Biochemistry and Biophysics 290, 543–548.

Hart, G.W., Slawson, C., Ramirez-Correa, G., and Lagerlof, O. (2011). Cross Talk Between O-GlcNAcylation and Phosphorylation: Roles in Signaling, Transcription, and Chronic Disease. Annual Review of Biochemistry 80, 825–858.

Hewagama, A., Gorelik, G., Patel, D., Liyanarachchi, P., Joseph McCune, W., Somers, E., Gonzalez-Rivera, T., The Michigan Lupus, C., Strickland, F., and Richardson, B. (2013). Overexpression of X-Linked genes in T cells from women with lupus. Journal of Autoimmunity 41, 60–71.

Ivanov, I.I., McKenzie, B.S., Zhou, L., Tadokoro, C.E., Lepelley, A., Lafaille, J.J., Cua, D.J., and Littman, D.R. (2006). The Orphan Nuclear Receptor RORγt Directs the Differentiation Program of Proinflammatory IL-17+ T Helper Cells. Cell 126, 1121–1133.

Kanneganti, T.-D., and Dixit, V.D. (2012). Immunological complications of obesity. Nature Immunology 13, 707.

Kidani, Y., and Bensinger, S.J. (2017). Reviewing the impact of lipid synthetic flux on Th17 function. Current Opinion in Immunology 46, 121–126.

Kreppel, L.K., and Hart, G.W. (1999). Regulation of a Cytosolic and Nuclear O-GlcNAc Transferase: ROLE OF THE TETRATRICOPEPTIDE REPEATS. Journal of Biological Chemistry 274, 32015–32022.

Lehman, D.M., Fu, D.-J., Freeman, A.B., Hunt, K.J., Leach, R.J., Johnson-Pais, T., Hamlington, J., Dyer, T.D., Arya, R., Abboud, H.,et al. (2005). A Single Nucleotide Polymorphism in MGEA5 Encoding O-GlcNAc-selective N-Acetyl-β-D-Glucosaminidase Is Associated With Type 2 Diabetes in Mexican Americans. Diabetes 54, 1214–1221.

Liu, R., Ma, X., Chen, L., Yang, Y., Zeng, Y., Gao, J., Jiang, W., Zhang, F., Li, D., Han, B.,et al. (2017). MicroRNA-15b Suppresses Th17 Differentiation and Is Associated with Pathogenesis of Multiple Sclerosis by Targeting O-GlcNAc Transferase. The Journal of Immunology.

Lund, P.J., Elias, J.E., and Davis, M.M. (2016). Global Analysis of O-GlcNAc Glycoproteins in Activated Human T Cells. The Journal of Immunology 197, 3086–3098.

Lydic, T.A., Busik, J.V., and Reid, G.E. (2014). A monophasic extraction strategy for the simultaneous lipidome analysis of polar and nonpolar retina lipids. Journal of Lipid Research 55, 1797–1809.

Lydic, T.A., Townsend, S., Adda, C.G., Collins, C., Mathivanan, S., and Reid, G.E. (2015). Rapid and comprehensive ‘shotgun’ lipidome profiling of colorectal cancer cell derived exosomes. Methods 87, 83–95.

Macintyre, A.N., Valerie A. Gerriets, Amanda G. Nichols, Ryan D. Michalek, Michael C. Rudolph, and Divino Deoliveira, S.M.A., E. Dale Abel, Benny J. Chen, Laura P. Hale, and Jeffrey C. Rathmell (2014). The Glucose Transporter Glut1 is Selectively Essential for CD4 T Cell Activation and Effector Function. Cell Metabolism 20, 61–72.

McDonald, J.G., Thompson, B.M., McCrum, E.C., and Russell, D.W. (2007). Extraction and Analysis of Sterols in Biological Matrices by High Performance Liquid Chromatography Electrospray Ionization Mass Spectrometry. In Methods in Enzymology (Academic Press), pp. 145–170.

McLaughlin, T., Liu, L.-F., Lamendola, C., Shen, L., Morton, J., Rivas, H., Winer, D., Tolentino, L., Choi, O., Zhang, H.,et al. (2014). T-Cell Profile in Adipose Tissue Is Associated With Insulin Resistance and Systemic Inflammation in Humans. Arteriosclerosis, Thrombosis, and Vascular Biology 34, 2637–2643.

Michalek, R.D., Gerriets, V.A., Jacobs, S.R., Macintyre, A.N., MacIver, N.J., Mason, E.F., Sullivan, S.A., Nichols, A.G., and Rathmell, J.C. (2011). Cutting Edge: Distinct Glycolytic and Lipid Oxidative Metabolic Programs Are Essential for Effector and Regulatory CD4+ T Cell Subsets. The Journal of Immunology 186, 3299–3303.

Mohammed, H., D’Santos, C., Serandour, Aurelien A., Ali, H.R., Brown, Gordon D., Atkins, A., Rueda, Oscar M., Holmes, Kelly A., Theodorou, V., Robinson, Jessica L.L.,et al. (2013). Endogenous Purification Reveals GREB1 as a Key Estrogen Receptor Regulatory Factor. Cell Reports 3, 342–349.

Ng, M., Fleming, T., Robinson, M., Thomson, B., Graetz, N., Margono, C., Mullany, E.C., Biryukov, S., Abbafati, C., Abera, S.F.,et al. (2014). Global, regional, and national prevalence of overweight and obesity in children and adults during 1980-2013: a systematic analysis for the Global Burden of Disease Study 2013. The Lancet 384, 766–781.

Ridker, P.M., Everett, B.M., Thuren, T., MacFadyen, J.G., Chang, W.H., Ballantyne, C., Fonseca, F., Nicolau, J., Koenig, W., Anker, S.D.,et al. (2017). Antiinflammatory Therapy with Canakinumab for Atherosclerotic Disease. New England Journal of Medicine 377, 1119–1131.

Santori, Fabio R., Huang, P., van de Pavert, Serge A., Douglass Jr, Eugene F., Leaver, David J., Haubrich, Brad A., Keber, R., Lorbek, G., Konijn, T., Rosales, Brittany N., et al. (2015). Identification of Natural RORγ Ligands that Regulate the Development of Lymphoid Cells. Cell Metabolism 21, 286–297.

Schindler, T.I., Wagner, J.-J., Goedicke-Fritz, S., Rogosch, T., Coccejus, V., Laudenbach, V., Nikolaizik, W., Härtel, C., Maier, R.F., Kerzel, S.,et al. (2017). TH17 Cell Frequency in Peripheral Blood Is Elevated in Overweight Children without Chronic Inflammatory Diseases. Frontiers in Immunology 8.

Schneider, C.A., Rasband, W.S., and Eliceiri, K.W. (2012). NIH Image to ImageJ: 25 years of Image Analysis. Nature methods 9, 671–675.

Shen, H., Goodall, J.C., and Hill Gaston, J.S. (2009). Frequency and phenotype of peripheral blood Th17 cells in ankylosing spondylitis and rheumatoid arthritis. Arthritis & Rheumatism 60, 1647–1656.

Shen, Y., and Tong, L. (2008). Structural evidence for direct interactions between the BRCT domains of human BRCA1 and a phospho-peptide from human ACC1. Biochemistry 47, 5767–5773.

Shen, Y., Wen, Z., Li, Y., Matteson, E.L., Hong, J., Goronzy, J.J., and Weyand, C.M. (2017). Metabolic control of the scaffold protein TKS5 in tissue-invasive, proinflammatory T cells. Nat Immunol 18, 1025–1034.

Slawson, C., Copeland, R.J., and Hart, G.W. (2010).O-GlcNAc signaling: a metabolic link between diabetes and cancer?. Trends in Biochemical Sciences 35, 547–555.

Soroosh, P., Wu, J., Xue, X., Song, J., Sutton, S.W., Sablad, M., Yu, J., Nelen, M.I., Liu, X., Castro, G.,et al. (2014). Oxysterols are agonist ligands of RORγt and drive Th17 cell differentiation. Proceedings of the National Academy of Sciences 111, 12163–12168.

Stadhouders, R., Lubberts, E., and Hendriks, R.W. (2018). A cellular and molecular view of T helper 17 cell plasticity in autoimmunity. Journal of Autoimmunity 87, 1–15.

Swamy, M., Pathak, S., Grzes, K.M., Damerow, S., Sinclair, L.V., van Aalten, D.M.F., and Cantrell, D.A. (2016). Glucose and glutamine fuel protein O-GlcNAcylation to control T cell self-renewal and malignancy. Nat Immunol 17, 712–720.

Wang, C., Yosef, N., Gaublomme, J., Wu, C., Lee, Y., Clish, Clary B., Kaminski, J., Xiao, S., Zu Horste, Gerd M., Pawlak, M.,et al. (2015). CD5L/AIM Regulates Lipid Biosynthesis and Restrains Th17 Cell Pathogenicity. Cell 163, 1413–1427.

Wang, X., Zhang, Y., Yang, Xuexian O., Nurieva, Roza I., Chang, Seon H., Ojeda, Sandra S., Kang, Hong S., Schluns, Kimberly S., Gui, J., Jetten, Anton M.,et al. (2012). Transcription of Il17 and Il17f Is Controlled by Conserved Noncoding Sequence 2. Immunity 36, 23–31.

Wei, J., and Tong, L. (2015). Crystal structure of the 500-kDa yeast acetyl-CoA carboxylase holoenzyme dimer. Nature 526, 723.

Winer, S., Paltser, G., Chan, Y., Tsui, H., Engleman, E., Winer, D., and Dosch, H.M. (2009). Obesity predisposes to Th17 bias. European Journal of Immunology 39, 2629–2635.

Yi, W., Clark, P.M., Mason, D.E., Keenan, M.C., Hill, C., Goddard, W.A., Peters, E.C., Driggers, E.M., and Hsieh-Wilson, L.C. (2012). Phosphofructokinase 1 Glycosylation Regulates Cell Growth and Metabolism. Science 337, 975–980.

Yuzwa, S.A., Macauley, M.S., Heinonen, J.E., Shan, X., Dennis, R.J., He, Y., Whitworth, G.E., Stubbs, K.A., McEachern, E.J., Davies, G.J.,et al. (2008). A potent mechanism-inspired O-GlcNAcase inhibitor that blocks phosphorylation of tau in vivo. Nat Chem Biol 4, 483–490.

Zhang, Z., Tan, E.P., VandenHull, N.J., Peterson, K.R., and Slawson, C. (2014). O-GlcNAcase Expression is Sensitive to Changes in O-GlcNAc Homeostasis. Frontiers in Endocrinology 5.

